# Quantifying visitor preferences for biodiversity: A community science approach to identify ecotourism assets

**DOI:** 10.1101/2025.05.26.656219

**Authors:** Keisuke Atsumi, Tatsuki Tsujino, Mao Nakaseko, Shougo Ogasawara, Ryosuke Tsuboi, Hiroki Taga, Takaaki Nishida

## Abstract

Ecotourism plays a vital role in conserving ecosystems and revitalising rural economies. Sustainable management requires strategies that effectively balance conservation needs with tourism promotion, necessitating a clear understanding of which landscapes and species attract visitors. However, identifying these “charismatic” assets has traditionally relied on costly and spatially limited surveys. This study leverages crowdsourced biodiversity data from digital platforms (iNaturalist and Biome) to identify nature-based recreation sites and visitor-preferred species in Inabe City, a rural Japanese municipality with high recreational visitation. We analysed 12,764 observations within the city and 238,625 reference observations from across Japan. By applying the Out-of-Area Activity Index (OAAI) to biological recording data, we distinguished visitors from residents with high accuracy. Spatial analyses using hurdle models revealed distinct preferences: while both groups frequented recreational facilities, residents concentrated on lowland forests, whereas visitors were significantly drawn to wetlands and mountain habitats. To quantify specific biological interests, we developed the Visitor Preference Index (VPI). This index highlighted that visitors disproportionately recorded aquatic insects (e.g., dragonflies) and endangered understory plants compared to their observations elsewhere, identifying these taxa as key ecological assets. These findings underscore the critical importance of managing highland forests and wetlands—habitats often vulnerable to degradation—to support both biodiversity and tourism. This study illustrates how crowdsourced biodiversity data can contribute to understanding human–nature interactions and offer a scalable tool for regional planning that integrates ecological conservation with sustainable tourism development.

## Introduction

Nature recreation is a fundamental cultural ecosystem service that enhances human well-being and contributes to community development (Millennium Ecosystem Assessment, 2005). People often travel across cities, regions, and countries to seek opportunities for nature experiences, bringing substantial economic benefits to areas with unique, well-preserved or well-managed ecosystems (Echeverri et al., 2022; Naidoo et al., 2016). Nature recreation can be motivated by landscapes and outdoor activities associated with specific places, as well as by the desire to observe particular species or ecosystems. Certain species—often termed charismatic species—disproportionately attract public attention or tourism interest (Albert et al., 2018; Lorimer, 2007; Veríssimo et al., 2009), although the identity and relevance of such species can vary across regions and visitor groups (Ressurreição et al., 2012; Tsuzuki et al., 2025). While ecotourism can offer economic opportunities, it also carries risks of environmental degradation, illegal collection of wildlife, and negative impacts on local communities through overtourism (Buckley, 2004; Buckley et al., 2016). Thus, regions aiming to revitalise their economies through nature-based tourism must balance tourism promotion with the protection of natural assets, including landscapes and species (Capocchi et al., 2019; Chun et al., 2020; Concu & Atzeni, 2012; Shannon et al., 2017). Achieving this balance requires an improved understanding of what people want to see and experience in nature, and where such preferences are expressed.

Traditional approaches for understanding recreational demand, such as questionnaires, interviews, or census-based statistics, are often costly and limited in spatial and temporal resolution, making it difficult to capture fine-scale human movement patterns (Chun et al., 2020). In recent years, social media data have increasingly been recognised as a valuable alternative source for analysing nature-based recreation, due to their large volume, broad spatial coverage, and relatively low cost (Ghermandi et al., 2023; Havinga et al., 2020). Because most social media content is generated via mobile devices, these data frequently include precise geolocation and timestamp information. As a result, platforms such as Flickr and Twitter have been widely used to map visitation patterns in protected areas (Chun et al., 2020; Levin et al., 2015), assess the attractiveness of landscapes or touristic sites (Lee et al., 2022; Martínez Pastur et al., 2016), and analyse large-scale tourism flows (Lenormand et al., 2018).

In analysing nature recreation patterns, distinguishing between local residents and visitors is important because these groups often differ in their motivations, familiarity with local environments, and ecological knowledge. Failing to separate residents from visitors can lead to biased interpretations of recreational demand, conflating local cultural use with tourism-driven visitation (Faggi et al., 2013; Vouligny et al., 2009). For this, geotagged social media data have also been used to infer users’ home places based on the density of posts or temporal frequency of posts. This residence-inference method, based on the assumption that individuals most frequently record activities near their home location, has become a widely adopted strategy for characterising mobility patterns from large-scale digital trace data (Ajao et al., 2015; Heikinheimo et al., 2022; Lenormand et al., 2018). Yet, existing applications have largely focused on where people go, rather than what biological components motivate their visits. As a result, the ecological drivers of visitation—particularly at the species level—remain poorly understood (Echeverri et al., 2022).

Beyond spatial visitation patterns, identifying the specific biological drivers—such as particular species or ecosystems—that attract people is essential for integrating conservation goals with tourism planning. Conservation culturomics studies have used Wikipedia page views to quantify species popularity at global scales (Mittermeier et al., 2021; Wong & Rosindell, 2022). However, such metrics often reflect broad public interest and media visibility, rather than the preferences of actual visitors to specific regions or the species that are locally present. A species that is popular online globally may not necessarily be the one driving visitation to a specific rural locale.

Biodiversity observation platforms, such as iNaturalist and Biome, offer a promising opportunity to address this gap. These platforms collect geotagged wildlife observations submitted by the public, providing fine-grained information on both the locations and taxonomic identities of observed species. Unlike passive online search data, these records represent active, in-situ engagement with nature. Consequently, they have been increasingly used in ecological and socio-environmental research, including studies of human–nature interactions and public engagement with biodiversity (Havinga et al., 2020; Silk et al., 2021; Stanford et al., 2024). Despite this potential, the use of biodiversity observation data to reveal species-level preferences of actual visitors remains underexplored.

Here, we extend residence-inference approaches originally developed for general social media platforms (Ajao et al., 2015; Heikinheimo et al., 2022; Lenormand et al., 2018) to biodiversity observation data, and combine them with species-level analyses of observation behaviour. Specifically, we propose a framework that (i) distinguishes visitors from residents based on users’ spatial posting patterns (Out-of-Area Activity Index: OAAI), and (ii) identifies species that are disproportionately recorded by visitors within a focal area relative to their observations elsewhere (Visitor Preference Index: VPI). This approach enables the detection of not only typical charismatic species but also “hidden” ecological assets that act as key drivers of nature-based visitation. To showcase the applicability of this framework, we analysed 251,389 biodiversity observations from the iNaturalist and Biome platforms across Japan, focusing on Inabe City as a case study. Inabe City is located on the outskirts of the Nagoya metropolitan area and is actively promoting ecotourism and green infrastructure initiatives. After validating our residence classification against independent questionnaire data, we compared spatial observation patterns between residents and visitors, examined environmental factors associated with these patterns, and identified species that disproportionately attract visitor attention. By linking human mobility and species-level interest, this study advances the use of crowdsourced biodiversity data for understanding nature-based recreation and provides practical insights for sustainable tourism planning and site management.

## Methods

### Study area

This study was conducted in Inabe City, Mie Prefecture, Japan (136.52°E, 35.16°N; Figure 1). The city covers 219.83 km^2^, characterized by a diverse topographical gradient from the Suzuka Mountains (max. 1,144 m) to lowland mosaics of residential and agricultural lands. Inabe serves as a critical peri-urban recreation hub for the Nagoya metropolitan area, attracting approximately 700,000 annual visitors.

**Figure 1.**
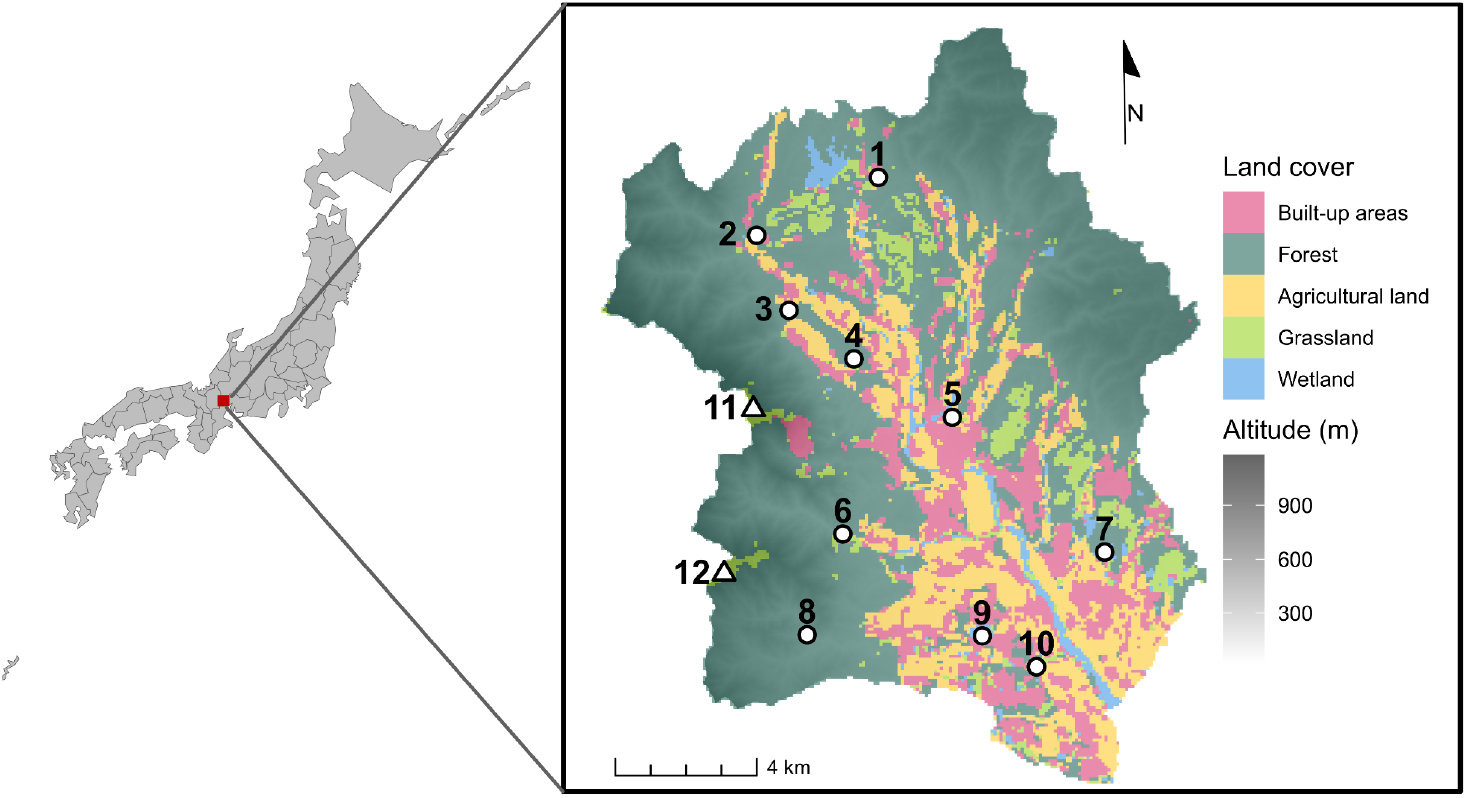
The map illustrates the land cover and topography of Inabe City, located in central Japan, and highlights key nature recreation sites. Numbered locations correspond to major parks, including (1) Inabe Agricultural Park, (2) Tatsuta Park, (3) Yane-no-nai-Gakko (a green infrastructure project for flood mitigation), (4) Furusato-no-Mori (an OECM site), (5) Nigiwai-no-Mori (a green-rich commercial complex), (6) Inabe Park, (7) Ryogaike Park, and (8) Nature Aquarium. It also shows campgrounds such as (9) Aogawakyo Camping Park and (10) Nordisk Hygge Circles UGAKEI, as well as prominent peaks, (11) Mt. Fujiwaradake and (12) Mt. Ryugatake.

The city is a particularly relevant site for studying nature-based solutions (NbS) and green infrastructure (GI) integration. Key GI sites include *Nigiwai-no-Mori*, an urban forest restoration project integrated with commercial functions, and *Yane-no-nai-gakko*, a multifunctional wetland restored for both biodiversity enhancement and flood mitigation. Furthermore, the *Furusato-no-Mori* site is recognized as a “Nationally Certified Sustainably Managed Natural Site” (an OECM-equivalent), reflecting the city’s commitment to balancing biodiversity conservation with public access. This established collaborative environment between municipal departments and research institutions provides an ideal platform for developing spatially explicit ecotourism strategies.

### Crowdsourced wildlife observation data

To capture high-resolution human-nature interactions, we utilized geotagged wildlife observations from two biodiversity community science platforms: iNaturalist and Biome. While iNaturalist is a globally recognized platform, Biome is a Japan-specific biodiversity observation application and widely used within the country for recording wildlife observations (Atsumi et al., 2024; Koide et al., 2023). Together, these platforms lower participation barriers through AI-assisted identification and community-based verification, providing an indirect indicator of public ecological interest.

Data was retrieved on 9 May 2025. To ensure high spatial precision—a prerequisite for landscape-scale planning—we applied strict filtering criteria: iNaturalist records were limited to those with a coordinate uncertainty below 200 m, and Biome records were restricted to observations with GPS-verified geotagged photos. Notably, for Biome users, we retrieved their entire observation history (both within and outside Inabe City) to enable a comparative analysis of recording behavior across different geographical contexts.

To analyze the spatial intensity of nature recreation, we calculated the number of unique users per 300-meter grid cell. This spatial resolution was selected to align with the typical scale of local recreational facilities and GI sites in the study area, which generally range from 200 to 500 meters. The use of unique user counts per cell, rather than total observations, mitigates bias from hyper-active contributors and provides a more accurate representation of spatial visitation patterns. Although the potential for individual users utilizing both platforms exists, the resulting double-counting is considered negligible given the substantial difference in user base sizes (see Results). All spatial analyses and statistical computations were performed using R version 4.4.1 (R Core Team, 2024).

### Resident-visitor classification and validation

To differentiate between residents and visitors among wildlife observers, we developed the Out-of-Area Activity Index (OAAI). The OAAI quantifies the proportion of an individual’s wildlife recording activity occurring outside the focal study area. We specifically calculated this index based on the number of unique observation days rather than the total volume of observations. Previous research on social media mobility analysis has demonstrated that temporal metrics (e.g., the number of active months) are more effective than post-count based methods for inferring home country of people visited a national park in South Africa, as they capture the persistence of presence while mitigating the bias of short-term “bursty” posting behavior typical of tourists (Heikinheimo et al., 2022). Given the ease of domestic mobility and the finer spatial scale of this study compared to cross-country analyses, we adopted “observation days” as the optimal temporal unit.

Accordingly, the OAAI is defined as the ratio of the number of days an individual recorded observations outside Inabe City to their total number of observation days across all locations. A high OAAI (approaching 1) reflects a transient presence in the city, typical of visitors whose primary activities occur elsewhere, whereas a low OAAI (approaching 0) suggests localized activity patterns characteristic of residents. Due to data availability, the OAAI was calculated specifically for users of the Biome platform, encompassing a national-scale dataset of 250,263 records. To ensure the robustness of the classification, we restricted the analysis to 387 users who had recorded observations on at least ten separate days.

The OAAI was validated against ground-truth data obtained from a voluntary survey integrated into a municipal biodiversity campaign, “Inabe Nature Positive Inacchu Quest,” conducted between 18 May and 30 November 2024. To ensure data quality, completion of the quest required recording at least three different wildlife species within the city. Upon meeting this requirement, users were presented with an optional, in-app questionnaire. Participants were explicitly informed that the survey was for academic research purposes and that their responses would be anonymized. Importantly, no monetary incentives were provided for responding, ensuring that participation was motivated by a genuine willingness to contribute to regional biodiversity efforts.

To differentiate between residents and visitors, we calculated the OAAI for each user. We validated this index using ground-truth data from the questionnaire survey, restricting the analysis to 44 respondents who met the activity threshold of ≥10 observation days. We employed logistic regression to assess the relationship between OAAI and actual residency status and determined optimal classification thresholds using the Youden Index via the *pROC* package (Robin et al., 2011). To ensure distinct separation between groups, we adopted conservative thresholds rather than a single cutoff (see Results 3.1). This research was conducted with the approval of the Research Ethics Committee of Kyoto Sangyo University.

### Spatial Patterns of Resident and Visitor Observations

We analysed the environmental factors (elevation and land cover, see Figure 1) associated with the observation patterns of residents and visitors. The highest elevation for each grid cell was extracted from the ALOS World 3D dataset provided by JAXA (Takaku et al., 2021), and standardised before statistical analyses. Land cover data in 2022 was obtained from the High-Resolution Land-Use and Land-Cover Map of Japan Version 23.12, also provided by JAXA (Hirayama et al., 2022). Land cover classifications were aggregated into five categories: (1) built, inherited from original classifications; (2) grass, inherited from original classifications; (3) forest, merging deciduous, evergreen, broadleaf and needleleaf forests, and bamboo; (4) agri, merging crop and rice categories; (5) wet, merging freshwater and wetland categories.

We employed Hurdle models to separately examine the number of resident and visitor observers in each grid cell, using pscl package (Zeileis et al., 2008). To eliminate the influence of inaccessible locations, we focused the analysis on 513 grid cells where at least one observer had recorded wildlife.

### Taxonomic patterns of observations by visitors

To identify species that specifically attract visitor attention within Inabe City, we developed the Visitor Preference Index (VPI). This index measures the extent to which a species is disproportionately recorded within the focal area relative to its national baseline. The VPI for species s is calculated as the difference between its local observation share and its overall observation share within the visitor group:

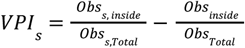

where *Obs* _*s, inside*_ is the number of observations of species *s* recorded within Inabe City, *Obs* _*s,Total*_ is the total number of observations of species *s* across all regions. The terms *Obs* _*inside*_ and *Obs*_*Total*_ represent the total number of visitor observations of all species within Inabe City and across all regions, respectively. The first term represents the proportion of observations of species s that occurred within the focal area, while the second term captures the baseline proportion of overall observation effort concentrated in Inabe City. By subtracting the latter from the former, the VPI isolates species-level disproportionality relative to general observation intensity. The VPI ranges from -1 to 1. A positive VPI indicates that the species is recorded in the focal area more frequently than expected based on its general recording frequency, suggesting local prominence that may reflect site-specific ecological characteristics, endemicity, or heightened recreational interest. Conversely, a VPI near or below zero suggests that the species’ presence in the area is consistent with or lower than its national average popularity among visitors. To ensure the reliability of the analysis and mitigate the influence of rare, stochastic observations, we calculated VPI and ranked species only for those observed by at least three different individuals within the city.

## Results

### Resident-visitor classification and validation

Logistic regression analysis confirmed a strong positive association between the OAAI and actual visitation status (Figure 2: *β*=10.00±5.12SE, *P*=0.05). While the Youden Index identified an optimal separation threshold of 0.45, we applied a more stringent classification to minimize error. Users with an OAAI < 0.2 were defined as residents, and those with an OAAI > 0.7 as visitors. This conservative approach provided a balanced accuracy of 0.95 and achieved 100% specificity for visitor identification, ensuring that residents were not misclassified as visitors (Figure 2).

**Figure 2.**
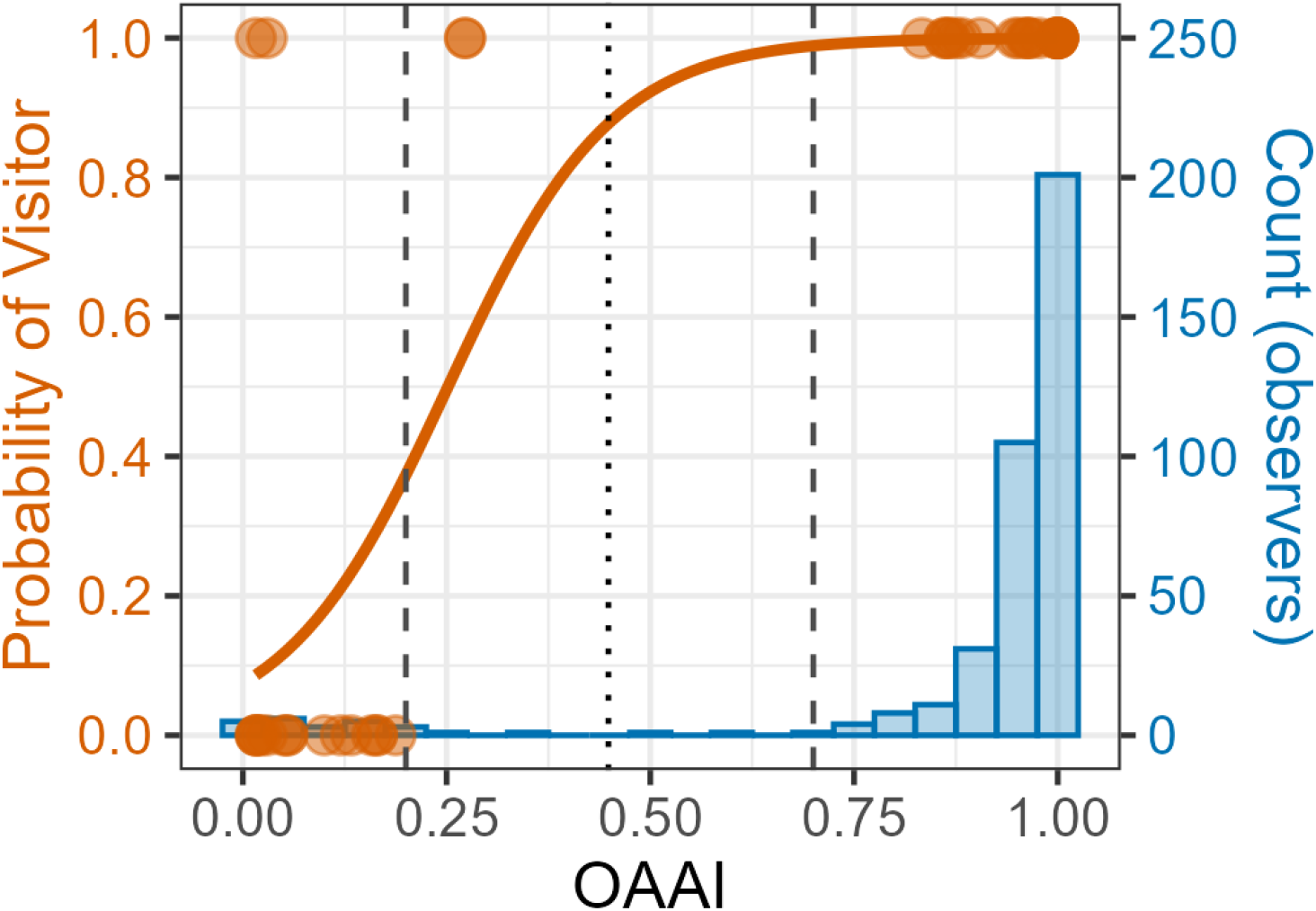
Validation of the Out-of-Area Activity Index (OAAI) against ground-truth data. The blue histogram shows the frequency distribution of users across the OAAI spectrum (right axis). Orange dots indicate the actual status of survey respondents (0: resident; 1: visitor), while the orange curve represents the logistic regression model predicting the probability of being a visitor (left axis). The black dotted vertical line marks the statistically optimal threshold. The dashed vertical lines denote the conservative thresholds adopted in this study: OAAI > 0.7 are classified as visitors, and OAAI < 0.2 as residents.

### Spatial Patterns of Resident and Visitor Observations

Biome and iNaturalist contained 11,638 and 1,126 wildlife observation records from 623 and 22 users, respectively, within Inabe City. Observation locations were broadly distributed across the city, except in the forested eastern region (Figure 3A). Outdoor recreation facilities, such as “Nigiwai-no-mori,” “Furusato-no-mori,” “Yane-no-nai-gakko,” and “Aogawakyo camping park,” saw significant wildlife observation activity (see also Figure 1 for recreational facilities). The highlands of the Suzuka mountains also attracted many observers.

**Figure 3.**
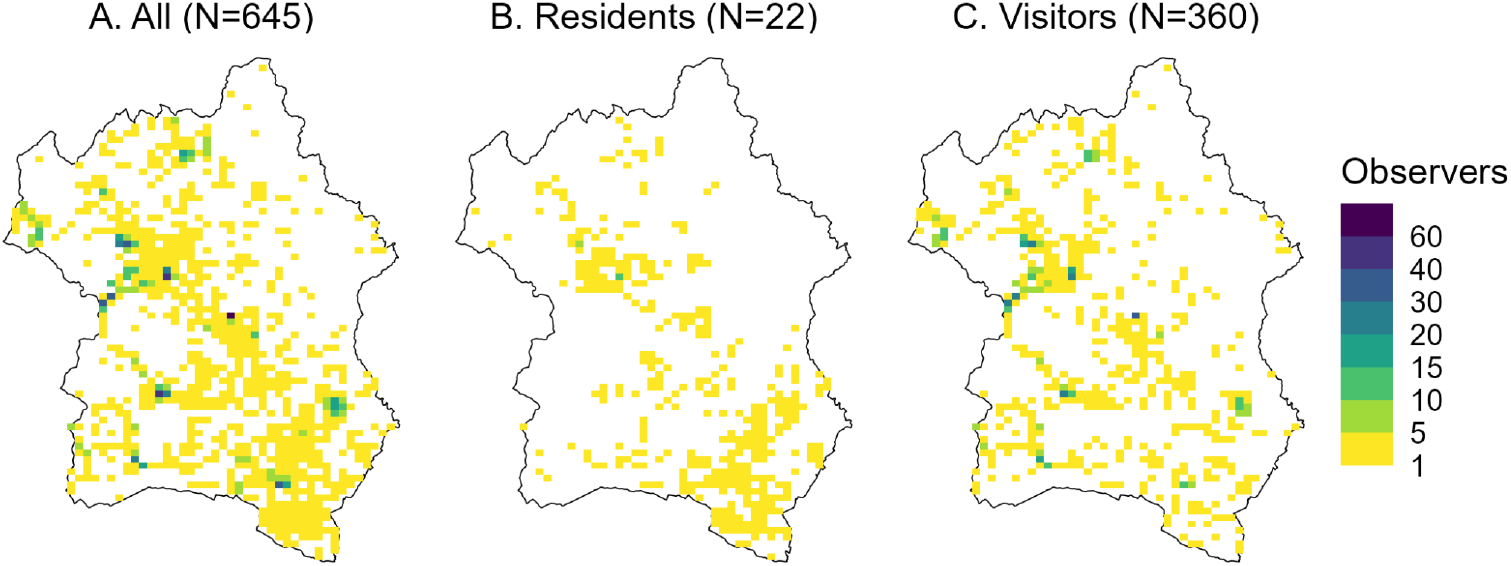
Spatial distribution of observers in Inabe City, displayed on a 300m resolution grid. Note that data for all observers were sourced from both iNaturalist and Biome, whereas residents and visitors were exclusively classified from Biome users based on their estimated residency status (OAAI).

Based on the OAAI, 22 observers were classified as residents and 360 as visitors. Spatial patterns of observations by residents and visitors showed weak negative correlation (number of observers in each grid cell: Spearman’s *ρ*=-0.15, *P*<0.001), suggesting spatial segregation between the two groups. Residents predominantly observed wildlife in the southern parts of the city, which are characterised by residential areas, croplands and industrial zones. They also frequented recreation facilities such as “Furusato-no-mori” and “Yane-no-Nai-Gakko” but not in mountain areas (Fig. 3B). In contrast, visitors avoided residential areas and focused their observations on recreation facilities such as “Nigiwai-no-mori,” “Furusato-no-mori”, camping sites, and Suzuka mountains (Fig. 3C).

Hurdle models further clarified these distinct spatial preferences by separating the probability of presence from the intensity of observation. Residents exhibited a strong preference for lowlands and a localized concentration of activity. Residents avoided highlands (elevation: *β*=-1.06±0.19, *P*<0.001), built-up areas (*β*=-2.52±1.16, *P*<0.05), and agricultural lands (*β*=-2.20±1.08, *P*<0.05). However, the count model revealed that in locations where observations did occur, the number of resident observers increased significantly in forests (*β*=3.08±0.90, *P*<0.001), wetlands (*β*=2.71±1.14, *P*<0.05), and built-up areas (*β*=2.52±0.93, *P*<0.01). This pattern implies that residents form observation hotspots in specific spaces in lowlands.

Visitors, in contrast, displayed a broad and active engagement with diverse environments. The zero hurdle model showed positive associations with almost all environmental variables, indicating a high probability of visitor presence across the landscape. Visitors were significantly more likely to make observations in highlands (elevation: *β*=0.99±0.32, *P*<0.01), forests (*β*=3.58±1.15, P<0.01), wetlands (*β*=3.28±1.58, *P*<0.05), built-up areas (*β*=2.81±1.11, *P*<0.05), and agricultural lands (*β*=2.04±1.03, *P*<0.05). The count model showed that the activity of visitors was further amplified by environmental features. Wetlands had the strongest effect on increasing visitor numbers (*β*=2.52±0.63, *P*<0.001), followed by grasslands (*β*=1.91±0.65, *P*<0.01), forests (*β*=1.53±0.50, *P*<0.01) and elevation (*β*=0.11±0.04, *P*<0.01). These results indicate that visitors explore a wide range of land covers, particularly nature-rich areas.

### Species-level visitor preferences identified by VPI

The analysis suggests a distinct visitor preference for specific taxa, identifying them as potential ecological assets for tourism in Inabe. The top 20 species with the highest VPI scores were predominantly aquatic insects and plants (Table 1). All exhibited positive index values, indicating that they were recorded by visitors more frequently than expected based on the national baseline.

**Table 1.**
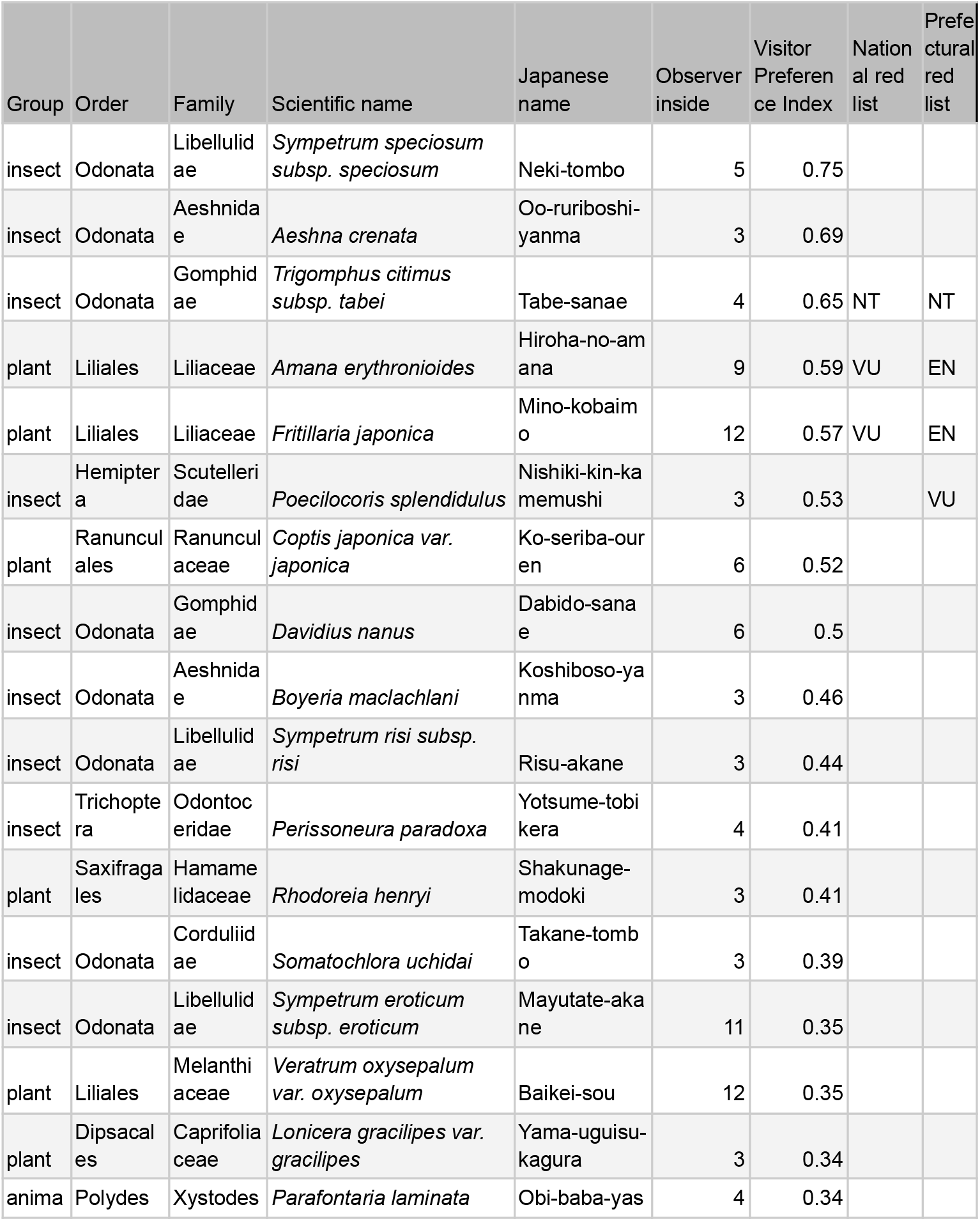
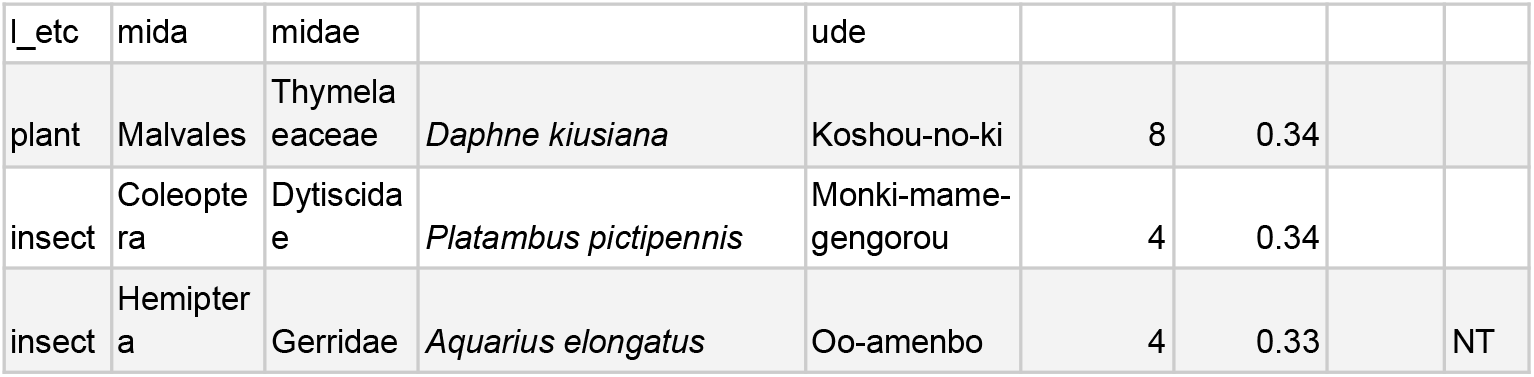
Top 20 species ranked by the Visitor Preference Index (VPI), excluding those recorded by fewer than three individuals in Inabe City. Species group classifications follow Atsumi et al. (2024). “Observers” refer to the number of individuals who recorded the species within the city. “National red list” and “prefectural red list” indicate the species’ conservation status on the Japan national and Mie Prefecture Red Lists. For the complete species list, see Table S1.

Aquatic insects appeared to be a key driver of visitor observations. This group included eight dragonfly species (Odonata), the diving beetle *Platambus pictipennis*, the caddisfly *Perissoneura paradoxa*, and the water strider *Aquarius elongatus*, which is listed as Near Threatened on the Mie Prefecture Red List. Native plants also featured prominently in the high-VPI list. The fourth and fifth-ranked species were endangered plants (*Amana erythronioides* and *Fritillaria japonica*: listed as Vulnerable on Japan’s national red list), naturally found in the Suzuka Mountains and are also ex-situ conserved in Furusato-no-mori. Other mountain-dwelling plants, such as *Coptis japonica var. japonica, Veratrum oxysepalum var. oxysepalum, Lonicera gracilipes var. gracilipes* and *Daphne kiusiana*, also appeared as top-rankers. These findings imply that visitors may be selectively targeting rare or characteristic biodiversity elements of the region.

## Discussion

### Distinct spatial behaviors of residents and visitors

Our analysis using the OAAI revealed that the majority of wildlife observations on digital platforms in Inabe City were contributed by visitors rather than residents. This dominance of visitors aligns with the city’s tourism statistics, where the annual number of visitors significantly exceeds the local population (ca. 700,000 vs. 44,000). While the OAAI achieved high precision in identifying visitors, the classification of residents included some uncertainty and a smaller sample size. Therefore, interpretations regarding resident behaviors require caution. Nevertheless, the results clearly delineated distinct spatial patterns between the two groups. Residents and visitors exhibited a “spatial segregation” in general observation activities. Residents tended to record wildlife in lowlands and areas with higher human impact, reflecting daily interactions with nearby nature. In contrast, visitors concentrated on specific natural assets, such as high-altitude trails in the Suzuka Mountains and freshwater environments such as wetlands. However, a notable “spatial overlap” was observed at specific Green Infrastructure (GI) sites. Facilities like “Furusato-no-Mori” (an OECM site) and “Yane-no-nai-gakko” (a flood mitigation basin) attracted observations from both groups (Fig. 1 and 3). This suggests that well-managed GI facilities are successfully functioning as nodes that connect local communities with tourism flows, fostering interactions between residents and visitors.

### Implications for ecotourism and conservation planning

The spatial preferences of visitors—favouring wetlands and mountains—strongly corroborated the findings from the VPI. A particularly notable discovery was the high visitor preference for aquatic insects (e.g., dragonflies, diving beetles), alongside the expected interest in rare mountain flora. While the aesthetic value of the region’s flora has been recognized in local conservation contexts, the potential of aquatic insects as “charismatic” tourism resources has been largely overlooked. The coherence between *where* visitors go (wetlands and mountains) and *what* they record (aquatic insects and understorey plants) suggests that these ecosystems are not merely background scenery but critical drivers of the visitor experience.

While it remains unclear whether these species are the primary motivation for visits or incidental discoveries during activities like hiking, the high VPI scores imply that specific taxa contribute significantly to the quality of the visitor experience. Species distribution models for these organisms may uncover “hidden gem sites” for tourism development (Kass et al., 2023).

Our results suggest that, to safeguard these ecotourism assets, conservation strategies should prioritize wetlands and mountain forests. In warm regions of Japan, wetlands are particularly vulnerable to rapid vegetation succession and drying, requiring active human intervention to maintain ecosystems (Tomita, 2014). Since many wetlands in Inabe remain unmanaged or inaccessible, active management to maintain open water surfaces—combined with improved accessibility—could enhance their value as tourism resources. In the Suzuka Mountains, increasing sika deer (*Cervus nippon*) populations pose a serious threat to the understory vegetation and natural grasslands (Uehira and Nishida, in prep). Therefore, active wildlife management, such as the installation of deer exclosures and population control, is crucial to maintain the quality of these botanical assets.

Conversely, the Yoro Mountains received far fewer observations compared to the Suzuka Mountains, likely due to limited hiking infrastructure and lower habitat suitability for popular plant species (Biome Inc., unpublished data). This indicates that tourism development strategies should respect the current zoning, focusing on the Suzuka range for recreation while maintaining the Yoro range primarily for forestry. Collaborations with destination management organisations (e.g., Green Creative Inabe) and local environmental agencies (e.g., Mt. Fujiwaradake Natural Science Museum) are essential to leverage these “hidden gem” assets while mitigating risks of overtourism. Highlighting biodiversity “assets” through the VPI can guide the development of regenerative tourism programs that not only utilize but also contribute to the conservation of these habitats.

### Utility of community science indices for landscape planning

This study demonstrated the utility of two indices, OAAI and VPI, for extracting actionable insights from unstructured community science data. The OAAI successfully adapted the logic of home-location inference from geotagged social media photos (Heikinheimo et al., 2022; Lenormand et al., 2018) to biological recording platforms. Its high specificity (100%) in identifying visitors allows planners to isolate “tourism demand” from “local usage” without relying on costly traditional surveys. In urban contexts, the index might misclassify individuals commuting for work or school as residents rather than true inhabitants. Nevertheless, even in such cases, the OAAI is likely to remain effective in distinguishing individuals who regularly engage with a given ecosystem from those whose interactions are limited to occasional visits.

The VPI was designed as a communication tool for local stakeholders. We deliberately calculated VPI based on the *number of records* rather than the *number of observers*. This approach captures how “readily” a species can be encountered and photographed in the region, which is a practical metric for tourism promotion. If the index were based solely on the number of observers, it might fail to highlight species that are locally abundant and frequently enjoyed by visitors but are relatively rare or difficult to observe outside the city. While this method carries a risk of inflation by a few enthusiastic users recording the same species repeatedly, we mitigated this by setting a minimum threshold for the number of observers (Table 1). In our dialogues with local stakeholders, this simple, record-based metric proved intuitive and effective for consensus building regarding which species to feature in tourism narratives.

The practical utility of the VPI is already being demonstrated in Inabe City. The visualization of “hidden” assets, such as aquatic insects, has provided a new common language for stakeholders who previously focused primarily on charismatic mammals or landscapes. Based on these findings, we have initiated dialogues with local government officials and tourism operators to design “regenerative tourism” programs. These programs aim to direct visitor interest toward these specific taxa while channeling tourism revenues into the conservation of their habitats (e.g., managing wetland vegetation).

## Conclusions

By treating biodiversity records accumulated on digital platforms as data reflecting human–nature interactions, this study developed a cost-effective and scalable approach to identify species attractive for visitors and nature observation hotspots. Leveraging crowdsourced biodiversity observations, we inferred which locations and species in Inabe City attract attention from both residents and visitors. While the popularity of green infrastructure facilities and rare or endemic plant species was anticipated, the unexpectedly high interest in aquatic insects highlights overlooked assets with ecotourism potential. These findings offer valuable insights to support collaboration among stakeholders, enabling more strategic and sustainable planning of ecotourism initiatives. In turn, such efforts can contribute to regional revitalisation by linking biodiversity conservation with local economic development.

## Acknowledgments

We thank Tsukasa Katayama and Manami Araki for their valuable feedback on the analyses, and Kenta Uchida for his comments on an earlier version of the manuscript. We are also grateful to all participants in the online survey. This work was supported by the Council for Science, Technology and Innovation (CSTI), Cross-ministerial Strategic Innovation Promotion Program (SIP), the 3rd period of SIP “Smart Infrastructure Management System” Grant Number JPJ012187 (Funding agency: Public Works Research Institute).

## Data availability

All code used in this study will be made publicly available on GitHub upon acceptance. A derived dataset, including aggregated observation counts, species attributes, and anonymized spatial units, will be shared via the Open Science Framework (OSF).

To protect observer privacy and prevent potential harm to wildlife, raw geolocated observation records from the biodiversity platforms will not be made publicly available. This is particularly important because some observations include photographs taken on private property, and disclosure of precise locations of threatened species may increase the risk of illegal collection.

For researchers seeking to replicate or extend this study, access to raw geolocated data may be granted upon reasonable request, subject to data use agreements and ethical approval, and only where such sharing does not conflict with privacy protection or conservation considerations.

## Declaration of generative AI and AI-assisted technologies in the manuscript preparation process

During the preparation of this work the authors used Gemini and ChatGPT in order to prepare R scripts and language editing. After using these tools, the authors reviewed and edited the content as needed and take full responsibility for the content of the published article.

## Notes

### Competing Interest Statement

Keisuke Atsumi and Hiroki Taga are employed by BIome Inc.; Shougo Ogasawara is employed by Pacific Consultants. The other authors have no conflicts of interest to disclose. None of the authors receive direct financial benefit from this study.

### Summary of Updates

While the concept remains unchanged, whole manuscript was revised.

