## Supplementary material for "Quantifying visitor preferences for biodiversity: A community science approach to identify ecotourism assets": Table S1: 4_taxon_visitor_ratio.html

Species particularly popular among Inabe visitors


Code 

- Show All Code
- Hide All Code

### Species particularly popular among Inabe visitors

```
pacman::p_load(
  tidyverse, data.table,
  plotly,reactable,
  kableExtra,
  knitr       # rmarkdown
  )

opts_chunk$set(
  prompt=TRUE, message=FALSE, comment="", warning=FALSE, echo=FALSE
  ) 
options(
  knitr.kable.NA = '',
  scipen=100           # do not show numbers using exponential
  )

conflicted::conflict_prefer_all('dplyr', quiet = TRUE)
conflicted::conflict_prefer_all('tidyr', quiet = TRUE)

dir_nas <- "//biome-synology/data_science/"

t <- fread("data/taxon/taxon_meta.csv") |> distinct() %>% 
  mutate(
    across(
      c(nationalRL, starts_with("prefRL_")),
      \(x) str_remove_all(x, ".*\\p{Han}")
  ))
u <- fread("data/user/oaai.csv")
```

### Compare inside/outside Inabe in visitors

Figure S1 shows the Visitor Preference Index and the number of
observers inside Inabe city.

##### Table S1

Table S1 summarises taxonomic information of each species, the number
of records inside the city, the number of observers inside the city, the
Visitor Preference Index and the status on Japan national and Mie
prefectural redl ists
